# Phage-encoded factor stimulates DNA degradation by the Hna anti-phage defense system

**DOI:** 10.1101/2025.11.12.688083

**Authors:** Matthew M. Hooper, Benjamin T. Hoover, Hongshan Zhang, Adam S. Franco, Ilya J. Finkelstein, David W. Taylor

## Abstract

Prokaryotic organisms have evolved unique strategies to acquire immunity against the constant threat of bacteriophage (phage) and mobile genetic elements. Hna is a broadly distributed anti-phage immune system that confers resistance against diverse phage by eliciting an abortive infection response. Using a combination of biochemistry, cryo-electron microscopy, and single-molecule fluorescence imaging, we reveal that Hna functions as a 3’—5’ single-stranded DNA exonuclease that forms an auto-inhibited dimer under physiological ATP concentrations. We observed that Hna autoinhibition can be overcome by incorporation of a phage-encoded single-stranded DNA binding protein (SSB), stimulating unregulated Hna nuclease activity. Furthermore, phage escape mutants encode SSB variants that evade Hna surveillance by adopting higher order stoichiometries with enhanced DNA binding affinity. Our work establishes the molecular basis of Hna-mediated anti-phage activity and provides novel insights into how phage-encoded proteins can directly stimulate a bacterial immune response.

## Introduction

Prokaryotic life is continually challenged by parasitic genetic elements, such as bacteriophage (phage), that contribute disproportionately to bacterial death and are ubiquitous to every environment in which bacteria can grow^1,2^. The environmental pressures imposed by phage have driven bacteria to evolve a diverse repertoire of resistance strategies to neutralize or limit the spread of infection^3,4,5,6^. Many well-characterized anti-phage systems, such as CRISPR-Cas and restriction modification (RM) systems, employ targeted nucleic acid degradation to disrupt replication of the phage genome^5,7,8^. By contrast, a distinct subclass of bacterial immune systems, known as abortive infection (abi) systems, often interfere with viral propagation by weaponizing host cellular processes and triggering cell death or dormancy to benefit the surrounding bacterial population^9,10,11^. Recent studies highlight the mechanistically diverse and underexplored strategies of abi systems, emphasizing the potential for the discovery of novel approaches used by bacteria to combat phage^12,13,14,15,16^.

Hna (helicase-nuclease abortive infection) is an abi defense system originally discovered in the gram-negative bacterium, *Sinorhizobium meliloti*, that has been identified in ∼2% of sequenced bacterial genomes^15,17,18,19^. Foundational work by Sather et al. established that Hna functions as a single-effector system to confer immunity against diverse families of tailed viruses (*Caudoviricetes*). Hna interferes with phage genome replication without releasing phage progeny via an unknown mechanism^15^. Hna possesses highly conserved N-terminal superfamily II (SF2) helicase motifs and a C-terminal PD-(D/E)XK nuclease domain. The cooperativity of nuclease and helicase modules is an established approach for defense against phage attack and has been characterized in functionally distinct systems such as Hachiman, Gabija, Nhi, and Cas3 ^14, 20, 21, 22, 23, 24, 25^. Conservation of both nuclease and helicase modules is often necessary for proper anti-phage activity, however, the interplay of each domain is highly variable across systems, allowing for unique modes of activation and regulation.

Recently, studies have identified several phage-encoded protein factors that induce abortive infection through direct or indirect activation of bacterial immune systems^26, 27, 28, 29^. Of growing interest are phage-encoded single-stranded DNA binding proteins (SSB) which have been shown to stimulate the anti-phage activity of several defense systems including Hachiman, Nhi, Retron-Eco8, AbpAB, and Hna^15,24,30,31^. Co-expression of a phage-encoded SSB is sufficient to induce Hna-mediated abi independent of phage infection^15^ and establishes a unique opportunity to explore the nature of Hna anti-phage activity and the external factors that regulate it.

In this study, we characterize the catalytic properties of the Hna system originating from *S. meliloti*. We demonstrate that Hna functions as a 3’—5’ exonuclease on single-stranded DNA and forms an auto-inhibitory homodimer upon binding ATP. Disrupting either the dimerization or ATP-binding domains restores Hna exonuclease activity, establishing a mode of self-regulation under normal cellular conditions. Addition of a phage-encoded SSB (5A SSB) stimulates Hna exonuclease activity even in the presence of ATP, disrupting Hna self-inhibition. Together, this work details the mechanism of Hna broad-spectrum anti-phage activity and contributes to our understanding of phage-encoded triggers for activation of abortive infection systems.

## Results

### Hna is a DNA exonuclease

To investigate the putative nuclease and helicase capabilities of Hna, we recombinantly expressed and purified the Hna gene originating from *Sinorhizobium meliloti* using *Escherichia coli*. Initial structural prediction using AlphaFold3 (AF3)^32^ suggested that Hna resembles a subclass of DinG-like proteins, termed ExoDinG, that exhibit 3’—5’ exonuclease activity on single-stranded DNA (ssDNA) but lack helicase activity^33,34^. Consistent with this prediction, we observed that Hna continually degrades ssDNA oligonucleotides in the presence of select divalent cations, Mg^2+^ and Mn^2+^ (**Fig. 1A, Extended Fig. 1A**). We then incorporated a series of nuclease-resistant phosphorothioate (PS) modifications at either the 3’ or 5’ end of the ssDNA to probe for directionality in nuclease activity. As expected, Hna was unable to degrade ssDNA with 3’ PS modifications but was unperturbed by the inclusion of 5’ PS modifications, supporting that Hna functions as a 3’—5’ exonuclease (**Fig. 1B, 1C**).

**Fig. 1.**
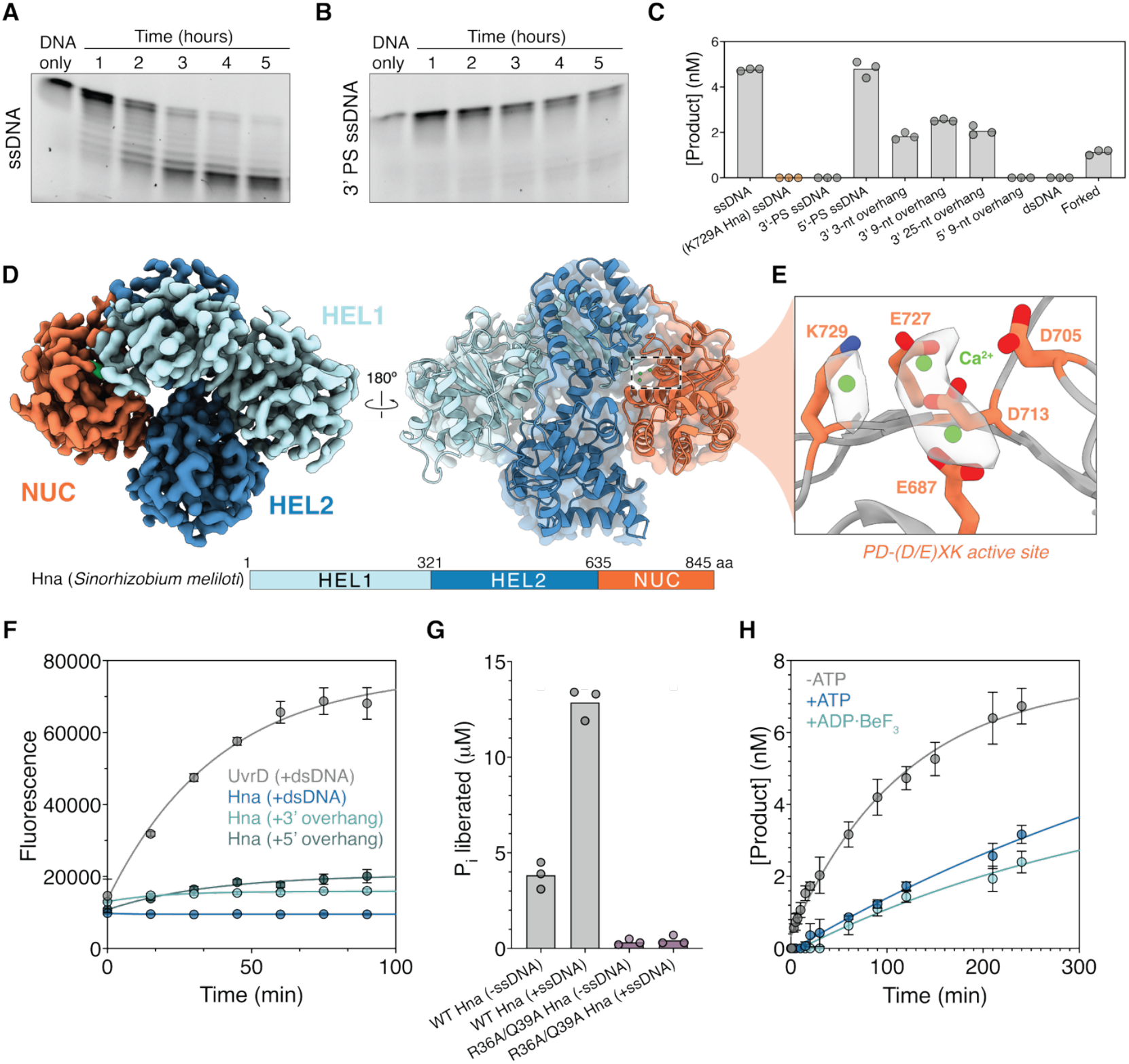
Hna is a 3’—5’ single-stranded DNA exonuclease that is negatively regulated by ATP. **A**, Exonuclease activity of Hna over time using fluorescently labeled, single-stranded DNA oligonucleotide without or **B**, with a series of 3’-phosphorothioate (PS) modifications. **C**, Quantification of DNA cleavage by Hna in the presence of various substrates using capillary electrophoresis. All reactions were incubated at 37°C for 1 hour. **D**, Cryo-EM structure of monomeric Hna at a resolution of 3.38 Å with Hna domain organization shown below (HEL1, Superfamily 2 helicase 1; HEL2, Superfamily 2 helicase 2; NUC, PD-(D/E)XK nuclease). **E**, Magnified view of PD-(D/E)XK nuclease active site showing significant catalytic residues and representative cryo-EM density for calcium ions (shown as green spheres). **F**, DNA unwinding on fully double-stranded or partially duplex substrates with 15-nucleotide overhangs on either the 3’ or 5’ end of the molecule. At double-stranded ends of each substrate, a 6-FAM label or Black Hole Quencher 1 (BHQ-1) modification is incorporated on either strand to quench fluorescence. All reactions were incubated at 37°C in the presence of ATP. The commercially available helicase, UvrD, included as a positive control. Data shown are mean ± standard deviation of three independent experiments. **G**, Malachite green ATPase assay using wild-type Hna or ATP-binding mutant (R36A/Q39A) with and without single-stranded DNA present. Phosphate concentration was determined after 30 minutes incubation. **H**, Quantification of ssDNA cleavage by Hna in the presence or absence of ATP or ADP·BeF3 using capillary electrophoresis. Data shown are the mean ± standard deviation of three independent experiments for each condition.

In addition to ssDNA, we also assessed whether Hna could degrade double-stranded DNA (dsDNA) or partially duplexed substrates. Using high-resolution capillary electrophoresis, we observed that Hna degrades all substrates that contain single-stranded 3’ ends and exhibits no nuclease activity with dsDNA or 5’ ssDNA overhangs (**Fig. 1C**). Hna cleavage products converge on lengths consistent with the length of ssDNA overhangs, suggesting that Hna likely degrades ssDNA processively but is unable to bypass double-stranded DNA junctions (**Extended Fig. 1C**).

To elucidate the molecular mechanism underlying Hna-mediated nuclease activity, we determined a cryo-electron microscopy (cryo-EM) structure of Hna at a resolution of 3.3 Å by combining Hna with 3’-PS ssDNA and the non-cleavage competent metal, Ca^2+^ (**Fig. 1D**). Hna is an 845-aa protein with two, N-terminal SF2 helicase domains (HEL1, HEL2) and a C-terminal PD-(D/E)XK nuclease-like domain (NUC)^35,36^. Comparison of Hna with structurally similar members of the DinG and XPD protein families revealed that Hna retains characteristic sequence motifs associated with ATP binding and hydrolysis in SF2 proteins, as well as a reduced Arch domain, yet lacks the canonical iron-sulfur cluster (**Extended Fig. 2**)^35, 37, 38, 39, 40^.

The C-terminal NUC domain of Hna contains the common catalytic core associated with PD-(D/E)XK nucleases (αβββαβ configuration) with critical residues coordinating three Ca^2+^ ions in the predicted active site (**Fig. 1E**)^36,41^. Mutation of a single residue (K729) was sufficient to completely abrogate Hna exonuclease activity (**Fig. 1C**). Despite the inclusion of ssDNA, we were unable to determine a DNA-bound structure of Hna. Lack of a stable, DNA-bound state suggests that Hna interacts only transiently with its ssDNA targets. This assumption is further supported by the observation that, under nuclease-active conditions, Hna exhibits exceptionally low binding affinity for ssDNA substrates (**Extended Fig. 3A**).

Next, we sought to characterize the potential DNA unwinding capabilities of Hna based on its highly conserved helicase domains. We incubated Hna with either fully dsDNA or partially duplexed DNA containing 3’ or 5’-overhangs in the presence of ATP. By incorporating a fluorophore on one DNA strand and a quencher in immediate proximity on the complementary strand, we were able to equate increased fluorescence to destabilization of the DNA duplex over time. Compared to the commercially available helicase, *Tte* UvrD, Hna exhibited no apparent DNA unwinding capabilities, regardless of the DNA substrate present (**Fig. 1F**)^42^. These findings confirm that, much like ExoDinG proteins, Hna functions as a 3’—5’ ssDNA exonuclease and does not possess DNA unwinding capabilities^33^.

Hna contains highly conserved SF2 helicase motifs yet exhibits no dsDNA unwinding functionality. We speculated that these motifs may instead use ATP binding and hydrolysis to modulate the exonuclease activity of Hna. Using a colorimetric-based assay, we first assessed the intrinsic and DNA-stimulated ability of Hna to hydrolyze ATP. These data revealed that Hna possesses innate ATPase activity that is significantly enhanced by the inclusion of ssDNA (**Fig. 1G, Extended Fig. 3E**). Mutation of two residues in the predicted ATP-binding site (R36A/Q39A), completely abrogated the ATPase activity of Hna (**Fig. 1G**)^35, 43^.

### ATP inhibits Hna nuclease activity

To recapitulate native levels of Hna nuclease activity, we performed a series of ssDNA cleavage assays using a wide range of ATP concentrations. Interestingly, we observed that, even at exceptionally low concentrations of ATP, Hna exonuclease activity was perturbed, indicating that ATP acts as a negative regulator of Hna nuclease activity (**Extended Fig. 3D**). Additionally, substitution of ATP for the nonhydrolyzable analog, ADP·BeF_3_, yielded comparable inhibition of DNA degradation, suggesting that Hna enters an inhibited state upon ATP binding rather than following ATP hydrolysis (**Fig. 1H**)^44,45^. To understand the molecular mechanism of ATP-mediated nuclease inhibition, we performed a native gel shift assay after incubating Hna with a combination of ssDNA and various co-factors including Mg^2+^ and ATP. We observed that Hna stochastically forms a higher-order complex that is greatly enriched in the presence of ATP (**Fig. 2A**). Moreover, the inclusion of ssDNA reduced the ratio of the higher-order species with respect to monomeric Hna. Collectively, these observations demonstrate that ATP binding by Hna promotes formation of an auto-inhibited Hna dimer that can be destabilized by ssDNA-stimulated ATP depletion over time.

**Fig. 2.**
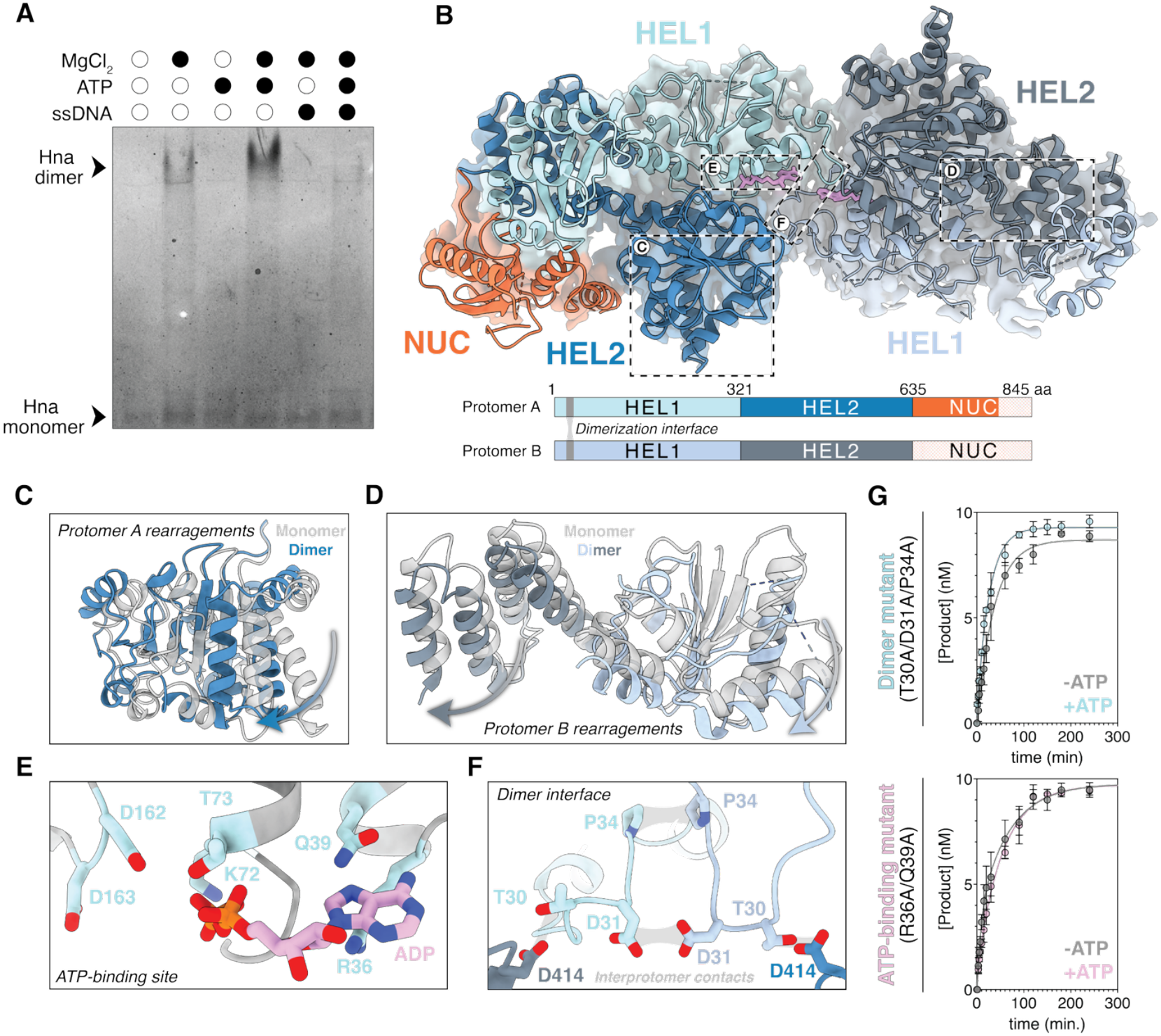
Hna forms an auto-inhibited dimer. **A**, Native gel shift assay to assess the basis of Hna dimerization. All reactions were incubated for 1 hour at 37°C then evaluated using non-denaturing gel electrophoresis. **B**, Cryo-EM structure of dimeric Hna at a resolution of 3.44 Å with domain organization of each protomer shown below. Dimerization interface highlighted in gray and unresolved portions of each nuclease domain represented by a crosshatched pattern. **C**, Magnified view of domain movements observed in protomer A or **D**, protomer B upon Hna dimerization. Structure of Hna dimer (shades of blue) overlaid with structure of Hna monomer (gray) with arrows depicting structural rearrangements. **E**, Magnified view of ATP-binding site highlighting functionally significant residues (Q motif: R36, Q39; Walker A motif: K72, T73; Walker B motif: D162, D163). ADP molecule occupying active site shown in pink. **F**, Magnified view of Hna dimer interface. Key residues in unstructured HEL1 loop (T30, D31, and P34) form interprotomer contacts (gray). **G**, Quantification of DNA cleavage by Hna mutants (Dimer mutant, T30A/D31A/P34A; ATP-binding mutant, R36A/Q39A) in the presence of ssDNA with or without the addition of ATP using capillary electrophoresis. Data shown are the mean ± standard deviation of three independent experiments for each condition.

Characteristic of SF2 helicase proteins is their ability to form stable homodimers, often with the dimeric state conferring increased activity^35, 46^. For Hna, however, we postulated that ATP-binding facilitates the formation of an Hna homodimer with reduced nuclease activity. To elucidate the molecular mechanism of Hna autoinhibition, we determined a cryo-EM structure of an Hna dimer in the presence of ADP·BeF_3_ and 3’-PS ssDNA at a global resolution of 3.4 Å (**Fig. 2B**). Hna assembles into a homodimer with psuedo-C2 symmetry via a highly conserved, unstructured loop in the HEL1 domain (**Fig. 2B, 2F**). The ATP-binding site in each Hna protomer was occupied by an ADP molecule stabilized by contacts from the highly conserved Q and Walker A motifs (**Fig. 2E, Extended Fig. 2B**). Two aspartate residues (D162 and D163) comprising the Walker B motif are also poised for coordination of a Mg^2+^ ion yet make no contacts due to the lack of the gamma phosphate group (**Fig. 2E**)^47^. Additionally, proximity of the ATPase active site to the dimerization interface provides a strong structural basis for stabilization of dimeric Hna upon ATP binding (**Fig. 2B**).

In comparison to our structure of an Hna monomer, Hna undergoes major conformational changes upon dimer formation. A cluster of centralized alpha helices in the HEL2 domain of protomer A undergoes an inward shift, enabling compaction of the two helicase domains and facilitating stabilization of the dimer interface (**Fig. 2C**). Critically, we observed that the NUC domain in both Hna protomers was almost entirely unresolved in the consensus structures, indicating a high degree of flexibility in this region. Rearrangement of both helicase domains in protomer B promotes flexibility of the NUC domain via a long, unstructured linker (**Fig. 2D**). These asymmetrical movements provide a physical basis for the regulation of Hna exonuclease activity through dimerization.

We aimed to elucidate the regulatory role of the Hna dimer by mutating key residues along the dimerization interface (T30A/D31A/P34A). DNA cleavage assays revealed that, compared to wild-type Hna, the Hna dimerization mutant exhibited significantly enhanced levels of ssDNA degradation that was unaffected by the inclusion of ATP. Additionally, we observed a similar phenotype when performing equivalent DNA cleavage experiments using the Hna ATP-binding mutant (**Fig. 2G**). These observations indicate that Hna dimer formation, stabilized through ATP binding, substantially inhibits Hna exonuclease activity. Consequently, disruption of either the Hna dimer interface or the ATP-binding domain leads to unregulated ssDNA exonuclease activity.

### Phage-encoded SSB activates Hna

Given the robust DNA-stimulated ATPase capabilities of Hna, we speculated that Hna may use ATP hydrolysis to translocate along ssDNA. To directly visualize Hna DNA interrogation we used single-molecule DNA curtains with Hna fluorescently labeled with ATTO647N, as previously described^51,52^. We first injected Hna into single-tethered ssDNA curtains in the absence of ATP. We observed very few, randomly distributed Hna binding events along the length of the ssDNA curtains. Upon introduction of ATP, Hna ssDNA binding was significantly enriched, establishing that Hna functions predominantly as an ATP-dependent single-stranded DNA binding protein (**Extended Fig. 3A, 3B**). Interestingly, in the absence of buffer flow, Hna rapidly diffuses along the DNA independent of ATP (**Extended Fig. 3C**). Lack of ATP-dependent translocase activity suggests that ATP binding assists in the localization of Hna to ssDNA targets but does not contribute to the innate ability of Hna to diffuse rapidly along lengths of ssDNA.

Previous work identified a predicted single-stranded DNA-binding protein (SSB) encoded by a T7-like podovirus, phage 5A, that elicits an Hna-mediated immune response even in the absence of true phage infection. Additionally, escape phage carrying missense mutations of the 5A SSB evaded detection by Hna^15^, implicating a potential interaction between Hna and the phage-encoded SSB leading to Hna-mediated abortive infection. Structural predictions revealed that the 5A SSB bears similarity to another SSB, gp2.5, encoded by bacteriophage T7^48, 49^. Based on this comparison, we suspected that the 5A SSB likely forms a stable dimer without DNA; however, highest likelihood predictions provided by AF3 suggest that Hna and the 5A SSB form a heterodimer composed of one copy of each protein (**Fig. 3B**)^32^. Formation of an Hna and 5A SSB complex would, therefore, necessitate destabilization of the Hna and 5A SSB dimers in favor of heterodimer assembly. To test this, we performed a native gel shift assay using Hna, 5A SSB, ssDNA, and ATP after pre-assembling each component into its dimeric form. We observed that a unique species forms in the presence of Hna, 5A SSB, and ATP, but not ssDNA, that is consistent in size with an Hna and 5A SSB heterodimer (**Fig. 3A**).

**Fig. 3.**
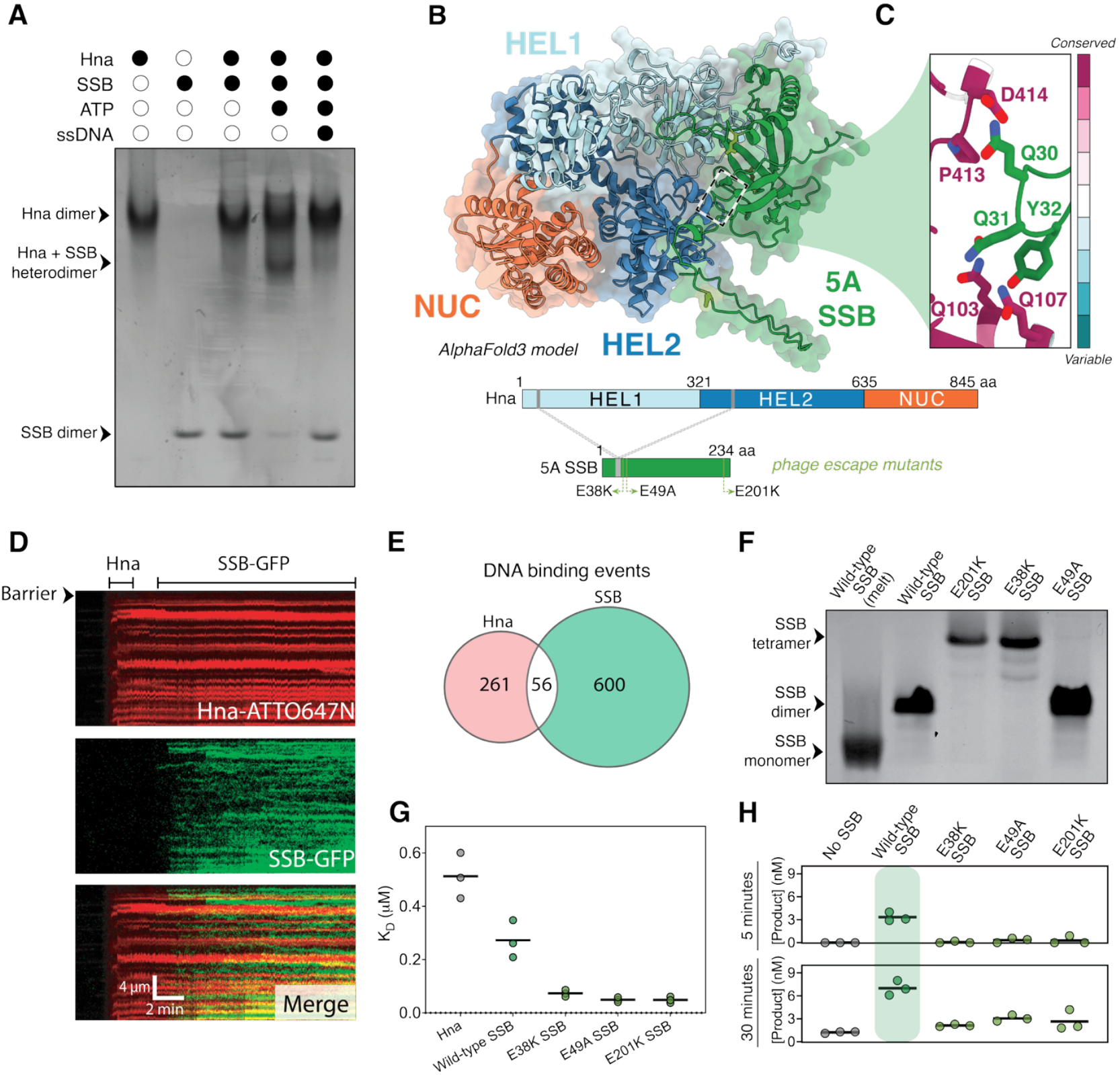
Phage-encoded 5A SSB stimulates Hna nuclease activity. **A**, Native gel shift assay to assess the basis of Hna and 5A SSB complex formation. Hna and 5A SSB allowed to dimerize independently then incubated at 37°C for 30 minutes in the presence of a combination of ssDNA and/or ATP. **B**, AlphaFold3 structural prediction of Hna and phage-encoded, 5A SSB heterodimer. Domain organization of each protein shown below with predicted protein-protein interface (gray) and characterized phage escape mutants (E38K, E49A, and E201K; green). **C**, Magnified view of predicted dimerization interface between Hna and 5A SSB. Hna residues are colored based on conservation analysis (ConSurf). **D**, Representative kymograph showing binding of ATTO647N-labeled Hna (red) and GFP-labeled 5A SSB (green) on single-stranded DNA curtains in the presence of ATP. Hna was injected first, followed by the 5A SSB to assess for colocalization events. **E**, Counts of Hna (red) and 5A SSB (green) binding events on ssDNA curtains. Overlap in binding events (white) indicates instances of colocalization. **F**, Native gel shift assay to evaluate the native stoichiometry of wild-type 5A SSB and 5A SSB escape mutants. A portion of the wild-type SSB reaction was melted at 95°C for 10 minutes to visualize the monomeric state. **G**, Binding affinity (KD) for ssDNA was determined for wild-type Hna and each 5A SSB variant using fluorescence anisotropy and 6-FAM labeled ssDNA oligonucleotide. Data shown are binding affinities determined from three independent binding curves, gray lines representing the mean of each condition tested. **H**, Quantification of DNA cleavage by wild-type Hna incubated with ATP and ssDNA with or without the addition of a 5A SSB variant using capillary electrophoresis. All reactions incubated at 37°C and evaluated at 5 and 30 minutes. Inclusion of wild-type 5A restores Hna nuclease activity in the presence of ATP at early time points (green rectangle). Data shown are concentration of product determined from three independent binding curves, gray lines representing the mean of each condition tested.

Surprisingly, Hna and the 5A SSB appeared to interact independent of ssDNA. To confirm this, we performed additional DNA curtains experiments, incorporating GFP-tagged 5A SSB (SSB-GFP) in addition to Hna using long ssDNA substrates. Although we observed extensive Hna and 5A SSB binding along the lengths of the DNA molecules, we did not record consistent or significant colocalization of the two proteins. This observation supports a model in which Hna and the 5A SSB form a complex prior to, or independent of, DNA binding (**Fig. 3D, 3E**).

Closer inspection of the AF3 model of the Hna and 5A SSB complex suggests that Hna and 5A SSB monomers compete for the same domain along the Hna dimerization interface (**Fig. 3C**). This provides a structural rationale for how the phage-encoded SSB may promote disassembly of the Hna homodimers in favor of Hna-5A SSB complex formation. Additionally, the proximity of the Hna ATP-binding site to the dimerization interface demonstrates why ATP binding is a shared prerequisite for both Hna homo- and heterodimer formation.

Given these observations, an intuitive strategy for evading detection by Hna would likely involve mutation of key residues responsible for Hna and 5A SSB dimer formation; however, upon mapping each of the previously studied 5A SSB escape mutants to the AF3 model, we noted that none of the mutated residues are within the dimerization interface (**Fig. 3B, 3C**). Due to each of the missense mutations conferring loss of a negatively charged residue, we speculated that the 5A SSB mutants may exhibit enhanced DNA binding that promotes localization to the phage DNA without being sequestered by Hna. We used fluorescence anisotropy to measure the binding affinity (K_D_) of Hna, wild-type 5A SSB, and three of the reported 5A SSB mutants (E38K, E49A, and E201K) for a single-stranded DNA substrate in the presence of ATP. As predicted, each of the 5A SSB mutants exhibited improved DNA binding compared to wild-type 5A SSB, with Hna possessing the weakest affinity for ssDNA even in the presence of ATP (**Fig. 3G, Extended Fig. 3A**). Additionally, native gel shift experiments revealed that two of the three 5A SSB mutants, E38K and E201K, adopt a unique, tetrameric stoichiometry in the absence of DNA as opposed to the expected dimeric state (**Fig. 3F**). Collectively, these data demonstrate that the 5A SSB mutants utilize diverse strategies to evade Hna detection through adopting higher order conformations that likely perturb formation of the Hna-SSB complex and increased affinity for ssDNA, further obstructing Hna-mediated anti-phage action.

Lastly, we wanted to assess how addition of 5A SSB affects Hna exonuclease activity. We evaluated the ability of Hna to degrade ssDNA in the presence of both ATP and either wild-type or mutant 5A SSB over time. Consistent with our results, inclusion of ATP substantially inhibited the exonuclease activity of Hna, however, upon the addition of 5A SSB, we observed substantial stimulation of ssDNA degradation by Hna (**Fig. 3H**). Conversely, no improvement in nuclease activity was observed with the addition of any of the 5A SSB mutants, suggesting that Hna remains in an ATP-dependent, autoinhibited state. Collectively, these data indicate that the phage-encoded 5A SSB is sufficient to activate Hna, allowing it to overcome autoinhibition and function as a constitutively active exonuclease during infection.

## Discussion

Hna is an abortive infection, bacterial immune system that provides protection from phage through a previously undetermined mechanism. Here, we show that Hna functions as a 3’—5’ ssDNA exonuclease that is negatively regulated through the formation of an auto-inhibited homodimer under physiologically relevant concentrations of ATP. Hna dimers can be destabilized through the incorporation of phage-derived SSBs, stimulating dysregulated nuclease activity that likely contributes to host cell death or dormancy. Overall, these data support that Hna is a broadly distributed anti-phage system that can surveil for phage-encoded SSB, resulting in constitutive exonuclease activation to confer resistance against phage infection.

We determined a cryo-EM structure of an Hna homodimer, revealing that the Hna dimer interface occurs between portions of the HEL1 domains of each subunit and encapsulates the ATP-binding site. At large, many well-characterized members of the SF2 helicase group do not exhibit nucleic acid unwinding properties^35, 46, 50^, thus, it was not surprising that Hna lacked DNA unwinding activity. It was, however, quite striking that Hna did not display any ATP-dependent translocase activity considering its highly conserved helicase machinery. Based on these observations, we speculate that the SF2 helicase motifs have largely been co-opted as self-regulating elements of Hna nuclease activity. Our observation that Hna readily forms an auto-inhibited dimer at physiologically-relevant concentrations of ATP supports the assumption that Hna exists in this inactive state during non-infection states. Due to low availability of ssDNA under non-stressed conditions^53, 54^, Hna likely remains unbound and trapped in a nuclease-incompetent state while acting as a surveillance system for phage-encoded factors, such as SSBs. Although we acknowledge that Hna possesses basal levels of intrinsic ATPase activity, this process is exceptionally slow and likely favors dimer reformation in the case of spontaneous dissolution due to ATP hydrolysis.

In addition to the highly conserved SF2 helicase domains, Hna has a functional C-terminal PD-(D/E)XK nuclease domain conferring 3’—5’ ssDNA exonuclease activity. Although our structure of an Hna monomer confirms expected organization of a PD-(D/E)XK nuclease motif^36, 41^, we note that the Hna NUC domain contains little to no structural or sequence similarity to other characterized nucleases outside of the conserved active site. This suggests that, while the catalytic residues are under strong selection, the surrounding scaffold has likely undergone extensive divergence like other characterized PD-(D/E)XK nuclease-containing anti-phage systems^36, 55, 56, 57^. We were unable to determine a DNA-bound state of Hna from our structural datasets despite the inclusion of ssDNA. This combined with the observation that Hna rapidly diffuses along ssDNA independent of ATP suggests that Hna has low affinity for ssDNA targets in the absence of ATP and likely exhibits a shallow DNA-binding mode, engaging primarily through surface contacts resulting in a minimal DNA footprint that is difficult to resolve using cryo-EM.

Despite the lack of resolvable ssDNA density, we observed that the nuclease active site was well-positioned for catalysis, with three calcium ions coordinated by the conserved PD-(D/E)XK residues. While PD-(D/E)XK nucleases often employ a two metal ion mechanism of DNA cleavage, several instances of three metal ion coordination have been reported within this family, speculated to provide additional structural support necessary for catalysis^58, 59, 60^. These observations support the assumption that stable docking of the NUC domain is required for Hna exonuclease activity and may be dependent on coordination of an additional metal ion. Domain rearrangements upon ATP binding and subsequent dimer formation may also disrupt or weaken metal ion binding within the NUC active site, providing an explanation for the lack of exonuclease activity and increased flexibility of the NUC domain in our Hna dimer structure.

Our work also provides direct biochemical evidence of Hna nuclease activation mediated by a phage-encoded SSB. Increased availability of ssDNA and production of 5A SSB during infection may promote Hna dimer dissolution through robust DNA-stimulated ATP hydrolysis and sequestration of Hna monomers by the SSBs (**Fig. 4**). We show that the 5A SSB allows Hna to overcome ATP-mediated nuclease inhibition and complexes with Hna independent of ssDNA. This observation supports previous work showing that Hna-mediated abortive infection can be induced exclusively by the inclusion of 5A SSB and without phage genomic material^15^. We speculate that SSB-induced activation of Hna nuclease activity may contribute to loss of host genome integrity or unregulated degradation of host plasmid, resulting in irreparable cell damage, and leading to abortive infection.

**Fig. 4.**
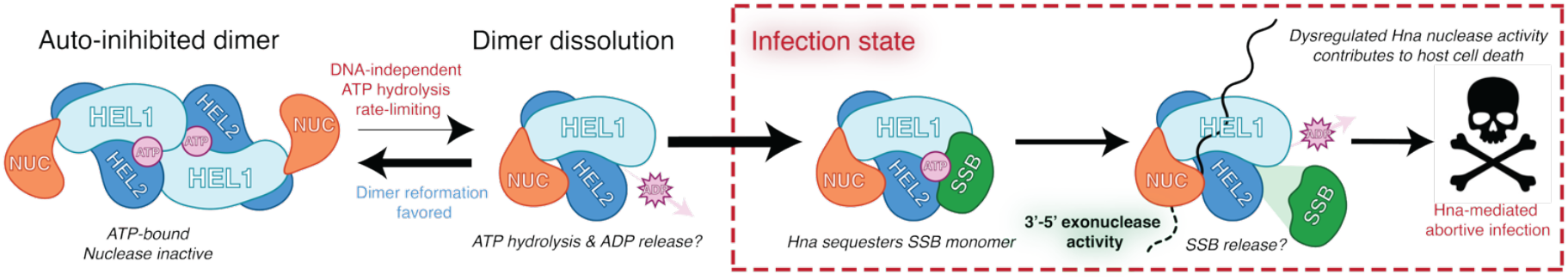
Proposed mechanism of Hna-mediated anti-phage action. Hna forms an auto-inhibited homodimer in the presence of ATP. Intrinsic ATPase activity contributing to dimer dissolution is likely slow and rate-limiting due to the low availability of single-stranded DNA during non-infection states. Upon phage infection, phage-encoded protein factors, such as single-stranded DNA binding proteins, can complex with Hna, releasing Hna from auto-inhibition, and stimulating Hna nuclease activity. Dysregulated DNA degradation by Hna during infection likely results in degradation of host plasmid or loss in host genome integrity, leading to abortive infection.

Hna functions uniquely as a single-effector ssDNA exonuclease with highly-conserved SF2 helicase domains involved in ATP hydrolysis and dimerization. We describe novel modes of Hna auto-inhibition and activation by a phage-encoded SSB, contributing to the expanding arsenal of known mechanisms employed by anti-phage systems to defend against infection. Future work is needed to resolve discrete modes of DNA-binding and catalysis by Hna. We anticipate that continued investigation of Hna orthologs will reveal diversity in function and recognition of phage-encoded factors across members of the Hna family of bacterial immune systems.

## Methods

### Plasmid construction

*S. meliloti* Hna and phage-encoded 5A SSB protein sequences as previously reported^15^ were reverse translated and codon optimized for expression in *E. coli*. The full-length Hna gene was commercially synthesized by Integrated DNA Technology (IDT) as a gBlock and assembled into a pET28 expression vector with a 6X N-terminal His tag, MBP tag, and TEV protease site using Gibson Assembly. Point mutations were introduced using site directed mutagenesis to generate each of the Hna variants. Each of the 5A SSB variants was commercially cloned into a pET28 expression vector with a 6X N-terminal His tag by Twist Biosciences.

The constructs described above were further modified for use in single-molecule fluorescence microscopy experiments. We ordered the coding sequence for a 3X-FLAG tag as a gBlock from IDT, then inserted it directly upstream of the Hna gene using Gibson Assembly. Additionally, we ordered a coding sequence for GFP as a gBlock from IDT and inserted it directly downstream of the 5A SSB gene using Gibson Assembly. These constructs were used to produce the Hna and SSB protein used for all single-molecule fluorescence microscopy experiments.

### Protein expression and purification

All Hna and 5A SSB proteins were recombinantly expressed and purified from *E. coli*. The expression plasmids described above were transformed into BL21 (DE3) or C41 (DE3) cells, plated on LB medium plates, and grown overnight at 37°C. Single colonies were used to inoculate 30 mL of LB media and cultured overnight at 37°C. For each liter of LB media, 10 mL of overnight starter culture was used to inoculate the expression cultures. Expression cultures were allowed to grow (37°C, 220 rpm) until reaching an optical density at 600 nm (OD_600_) of 0.6-0.7, then induced with 0.5 mM isopropyl-β-d-1-thiogalactopyranoside (IPTG) for 18 hours at 16°C (180 rpm).

Following induction, cells were pelleted by centrifugation and lysed via sonication in buffer containing 20 mM Tris-HCl (pH 7.5), 10% glycerol, 500 mM NaCl, 0.5 mM tris(2-carboxyethyl)phosphine (TCEP), 10 mM MgCl_2_, 1X DNase, and Pearce protease inhibitor tablets (Thermo Scientific). The cell lysate was clarified via ultracentrifugation and loaded onto a HisTrap HP column equilibrated in buffer containing 20 mM Tris-HCl (pH 7.5), 10% glycerol, 500 mM NaCl, and 0.5 mM TCEP, and eluted in buffer containing 20 mM Tris-HCl (pH 7.5), 10% glycerol, 500 mM NaCl, 0.5 mM TCEP, and 250 mM imidazole (pH 8). Fractions containing the target protein were added to Spectra/Por dialysis tubing (12-14 kD MWCO) and incubated in buffer containing 20 mM HEPES (pH 7.5), 10% glycerol, 150 mM KCl, and 0.5 mM TCEP at 4°C overnight. For Hna proteins, TEV protease was added for removal of affinity and MBP tags during dialysis. Dialyzed protein was concentrated using a 10 kDa or 50 kDa MWCO centrifugal filter (Sigma-Aldrich) and purified by size-exclusion chromatography (Superose 6 10/300; GE Healthcare) in buffer containing 20 mM HEPES (pH 7.5), 10% glycerol, 150 mM KCl, and 0.5 mM TCEP. Purified proteins were concentrated, aliquoted, flash-frozen using liquid nitrogen, and stored at −80°C.

### DNA cleavage assay

Hna cleavage assays were conducted using a buffer containing 20 mM Tris-HCl (pH 7.5), 100 mM KCl, 10 mM MgCl_2_, and 5% glycerol with 100 nM protein and 10 nM DNA. For select experiments, ATP or ADP·BeF_3_ were added at 1 mM unless otherwise specified. All DNA substrates tested are listed in Extended Table 1. Reactions were incubated at 37°C for the indicated time and terminated with the addition of 50 mM EDTA. For gel-based assays, Proteinase K and SDS were individually added to each reaction, mixed with TBE-Urea Sample Buffer (Novex), loaded onto a 15% TBE-Urea PAGE gel, and DNA products were allowed to separate for 2 hours at 90V. Gels were imaged using a fluorescence scanner. For analysis via capillary electrophoresis, reactions were combined with Hi-Di formamide (Applied Biosystems) and processed using a 3730 series DNA Analyzer (Applied Biosystems). Peaks corresponding to unique DNA species and sizes were quantified using the GeneMapper software.

### Cryo-EM sample preparation and data collection

To determine a structure of an Hna monomer, 15 µM Hna was incubated with 10 mM CaCl_2_ and 10 µM 3’-PS single-stranded DNA for 5 minutes at 37°C. 2.5 µL of sample was applied to glow discharged holey carbon grids (Quantifoil 1.2/1.3), blotted for 8 seconds with a blot force of 0, at 4°C and 100% humidity, then rapidly plunged into liquid ethane using an FEI Vitrobot Mark IV (Thermo Fisher). For the dimeric Hna structure, 15 µM Hna was incubated with 10 mM MgCl_2_, 10 µM 3’-PS ssDNA, and 1 mM ADP·BeF_3_ for 20 minutes at 37°C. 2.5 µL of sample was applied to glow discharged holey carbon grids (Quantifoil 1.2/1.3), blotted for 7 seconds with a blot force of 1, at 4°C and 100% humidity, then rapidly plunged into liquid ethane using an FEI Vitrobot Mark IV (Thermo Fisher). For both structures, data was collected using a FEI Glacios cryo-EM microscope (200 kV) equipped with a Falcon 4 direct electron detector (Gatan). Movies were recorded in SerialEM with a pixel size of 0.94 Å and a total exposure time of 15 seconds for an accumulated dose of 49 e^-^/Å^2^.

### Cryo-EM data processing and model building

All cryo-EM data processing was performed using CryoSPARC. Particles were initially picked by applying a minimum and maximum particle diameter of 70 and 100 Å, respectively for the monomeric dataset, and 70 and 150 Å, respectively for the dimeric dataset. Particle picks were manually inspected to eliminate particle outliers.

#### Hna monomer dataset

Particles were extracted with a box size of 300 pixels with a fourier crop to 128 pixels and then classified into 50 2D classes. 2D classes were manually selected then processed through one round of ab-initio volume reconstruction and heterogeneous refinement. The representative 3D class was further refined using non-uniform refinement. The resulting volume was used to generate 50, 2D templates that were used to guide particle picking via template picker. The refined set of particles was manually inspected and extracted with a box size of 284 pixels with a fourier crop to 128 pixels and then classified into 50 2D classes. 2D classes were manually selected then processed through two rounds of ab-initio volume reconstruction and heterogenous refinement with 5 classes for each. The representative 3D class from the final round of heterogenous refinement was further processed using non-uniform refinement, particles were unbinned, then underwent a final round of non-uniform refinement to determine a 3.38 Å structure of an Hna monomer.

#### Hna dimer dataset

Particles were extracted with a box size of 320 pixels with a fourier crop of 128 pixels and then classified into 50 2D classes. 2D classes were manually selected then processed through one round of ab-initio volume reconstruction and heterogeneous refinement. The representative 3D class was further refined using non-uniform refinement and then evaluated for particle heterogeneity using 3D classification. Volumes representing the same consensus structure were combined and refined using non-uniform refinement. Particles were unbinned and over-represented views were removed using the rebalance orientations job. The final set of particles was input into a reconstruct only job to determine a 3.44 Å structure of an Hna dimer.

Both Hna structures were initially rigid body fit using models generated from AlphaFold3 predictions within ChimeraX. Domain conformations were individually rigid body fit, then further refined through interactive rounds of modeling in Isolde and Coot. Models were subsequently processed using real space refinement in Phenix to generate final validation scores. Detailed structural analysis pipeline and validation statistics for both structures provided in Extended Figure 4 and Extended Table 2, respectively.

### DNA unwinding assay

Double-stranded DNA duplexes were prepared by combining either a 3’ or 5’-fluorescently labeled ssDNA oligonucleotide with a complementary ssDNA containing either a 3’ or 5’-Black Hole Quencher (BHQ-1) in a 1:2 molar ratio. Each mixture was heated to 95°C for 2 minutes and then slowly cooled at room temperature over the span of 4 hours. The DNA unwinding assay was performed by combining 100 nM of Hna with 10 nM of fully duplexed DNA or partially duplexed DNA containing either a 3’ or 5’ ssDNA overhang in a buffer containing 20 mM Tris-HCl (pH 7.5), 100 mM KCl, 10 mM MgCl_2_, 5% glycerol, and 1 mM ATP. Reactions were monitored in real time at 37°C for up to 90 minutes in the presence of an unlabeled strand identical to the FAM-labeled strand at 10X molar surplus to sequester the complementary strand upon DNA unwinding. Due to fluorescence quenching while duplexed, we monitored for increases in fluorescence over time using a CLARIOstar Plus plate reader (BMG Labtech).

### ATPase assay

Hna ATPase activity was assessed using a Malachite green phosphate assay kit (Sigma-Aldrich). Phosphate standards were prepared as detailed by the kit. Reactions were conducted using 100 nM Hna and a reaction buffer containing 20 mM Tris-HCl (pH 7.5), 100 mM KCl, 10 mM MgCl_2_, 5% glycerol, and 20 µM ATP, ADP·BeF_3_, AMP-PNP, ADP, CTP, or GTP with or without the inclusion of 10 nM unlabeled ssDNA. Each reaction was performed in triplicate and incubated at 37°C for 20 minutes. Reagent provided by the kit was added to each reaction, incubated at room temperature for 30 minutes, and then absorbance values at 620 nm were recorded for each reaction using a CLARIOstar Plus plate reader (BMG Labtech). We determined the concentration of free phosphate produced in each reaction by converting the raw fluorescence values to concentration using the standard curve generated.

### Electrophoretic mobility shift assay

To evaluate the basis of Hna dimerization, 1 µM Hna was pre-incubated for up to 2 hours at 37°C in a buffer containing 20 mM Tris-HCl (pH 7.5), 100 mM KCl, 10 mM MgCl_2_, and 5% glycerol with 10 nM ssDNA and/or 1 mM ATP. Reactions were loaded onto a non-denaturing 4-20% Tris-glycine protein gel (Invitrogen) and run at 100 V for 90 minutes at 4°C. For assessing Hna and SSB interaction and SSB stoichiometry, each protein was allowed to independently oligomerize for 1 hour prior to initiating the reaction. Hna and SSB were combined in equimolar concentration using the buffer composition described above in the presence or absence of 10 nM ssDNA and/or 1 mM ATP. Reactions were loaded onto a non-denaturing 4-20% Tris-glycine protein gel and run at 100V for 45-90 minutes at 4°C.

### Fluorescence anisotropy binding assay

Hna or 5A SSB were included at the highest concentration possible in a buffer containing 20 mM Tris-HCl (pH 7.5), 100 mM KCl, 10 mM MgCl_2_, 5% glycerol, and 0.05% Tween-20. We performed a series of 2-fold dilutions of each protein in triplicate, then combined each dilution with 10 nM of FAM-labeled single-stranded DNA with 3’-phosphorothioate modifications to observe binding without degradation. Each reaction was incubated for 30 minutes at 37°C in the presence or absence of ATP, then anisotropy values were recorded using a CLARIOstar Plus plate reader (BMG Labtech) equipped with a fluorescence polarization optical module. Data were fit using a one-step binding model to define the K_D_ for each unique protein tested.

### Single-molecule fluorescence microscopy

Single-molecule fluorescent images were collected using a customized prism TIRF microscopy-based inverted Nikon Ti-E microscope system equipped with a motorized stage (Prio ProScan II H117). The sample was illuminated with a 488-nm laser (Coherent Sapphire) and a 637-nm laser (Coherent OBIS) split by a 638-nm dichroic beam splitter (Chroma). Two color imaging was recorded using dual electron-multiplying charge-coupled device (EMCCD) cameras (Andor iXon DU897). Subsequent files were exported as uncompressed TIFF stacks and further analyzed in FIJI. Flowcells used for single-molecule DNA experiments were prepared as previously described51. All the single-molecule experiments were conducted in the imaging buffer (40 mM Tris-HCl pH 8.0, 50 mM NaCl, 2 mM MgCl2, 1 mM DTT and 0.2 mg mL-1 BSA) with or without 1 mM ATP at 37°C.

For the fluorescent labeling of Hna, protein was conjugated with monoclonal anti-Flag antibody (Sigma Aldrich, F1804) and anti-Mouse-IgG-ATTO647N antibody produced in goat (Sigma Aldrich, 50185) on ice for 10 minutes. The mixture was then diluted to a total volume of 150 μL imaging buffer. Immediately after the conjugation and dilution, the fluorescently labeled protein was injected into the flowcell at a 0.2 mL min-1 flow rate.

To image the binding of Hna on ssDNA and its colocalization with 5A SSB-GFP, the buffer flow was set to 0.2 mL min^-1^ flow rate. To observe Hna diffusion events, flow was temporarily turned off during imaging.

## Acknowledgements

This work was supported in part by Welch Foundation grant F-1938 (to D.W.T.) and the National Institutes of Health R35GM138348 (to D.W.T.). The content is solely the responsibility of the authors and does not necessarily represent the official views of the National Institutes of Health. We thank Kaoling Guan, Grace Hibshman, Jacquelyn Wright, and Nasim Abdi for insightful discussions and comments on the manuscript. Data were collected at the Sauer Structural Biology Laboratory at the University of Texas at Austin. We thank Axel Brilot and Evan Schwartz for assisting with cryo-EM data collection and processing.

## Author Contributions

The study was conceived by M.M.H. All protein purification and biochemical assays were performed by M.M.H. and B.T.H. Cloning was performed by M.M.H., B.T.H., and A.S.F. Cryo-EM data collection and structural analysis was performed by M.M.H. Single-molecule imaging and analysis performed by H.Z. Writing of the initial draft performed by M.M.H. with all authors reviewing, editing, and approving of the manuscript. Supervision and funding of the study was provided by I.J.F. and D.W.T.

## Data Availability

The structures and associated atomic coordinates have been deposited into the Electron Microscopy Data Bank (EMDB) and Protein Data Bank (PDB) with accession codes: Hna Monomer (EMD-72967 and PDB 9YHN) and Hna Dimer (EMD-73047 and PDB 9YKJ).

