## Extended Data for "Phage-encoded factor stimulates DNA degradation by the Hna anti-phage defense system"

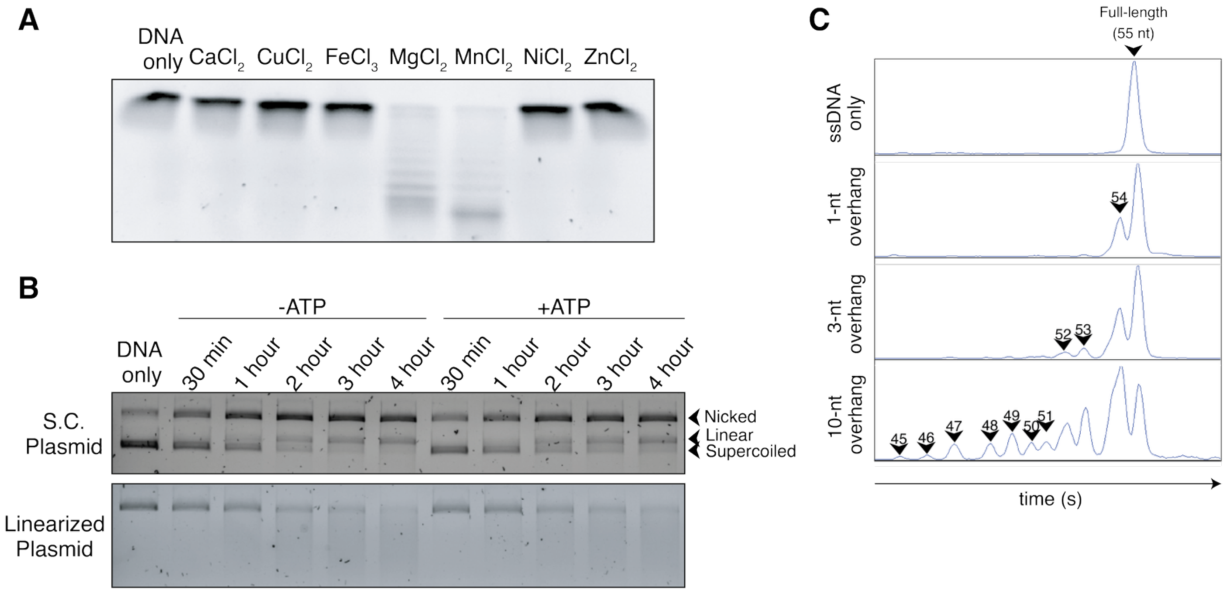


**Extended Data Fig. 1 | Hna degrades single-stranded and plasmid DNA. A,** Exonuclease activity of Hna over time using fluorescently labeled, single-stranded DNA oligonucleotide in the presence of various metals. **B**, Time course Hna cleaved using supercoiled (S.C.) or linearized plasmid DNA in the presence of absence of ATP. Distinct species of DNA (supercoiled, nicked, and linear) are denoted on the right. **C,** Raw fluorescence peaks corresponding to differently sized fluorescently-labeled cleavage products. DNA only negative control (55-nucleotides) shown at the top. Hna was incubated with each of the various 3'-overhang substrates (left) for 30 min. at 37°C.

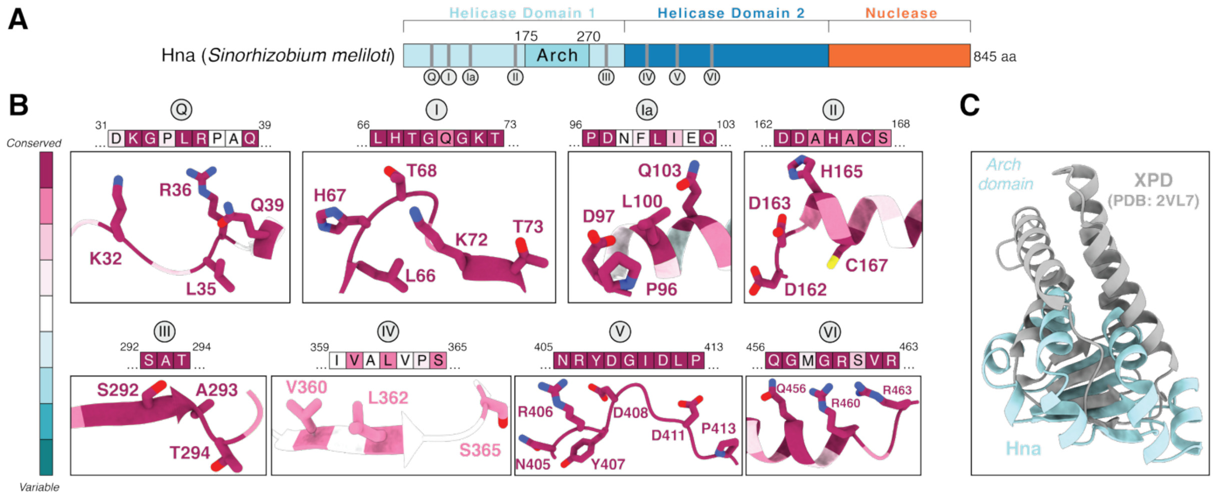


**Extended Data Fig. 2 | Hna possesses highly conserved SF2 helicase motifs. A,** Domain organization of Hna from *S. meliloti*. The two helicase modules and C-terminal nuclease domain are shown in blue and orange, respectively. Canonical HEL1 and HEL2 functional motifs are labeled, as well as the HEL1 Arch domain. **B,** Magnified view of each identified SF2 helicase motif. Residues are colored by conservation with position and sequence listed above. **C,** zoomed in view of Hna Arch domain (light blue) compared to XPD (PDB: 2VL7) Arch domain (gray) illustrating conserved core that is expanded in XPD compared to Hna

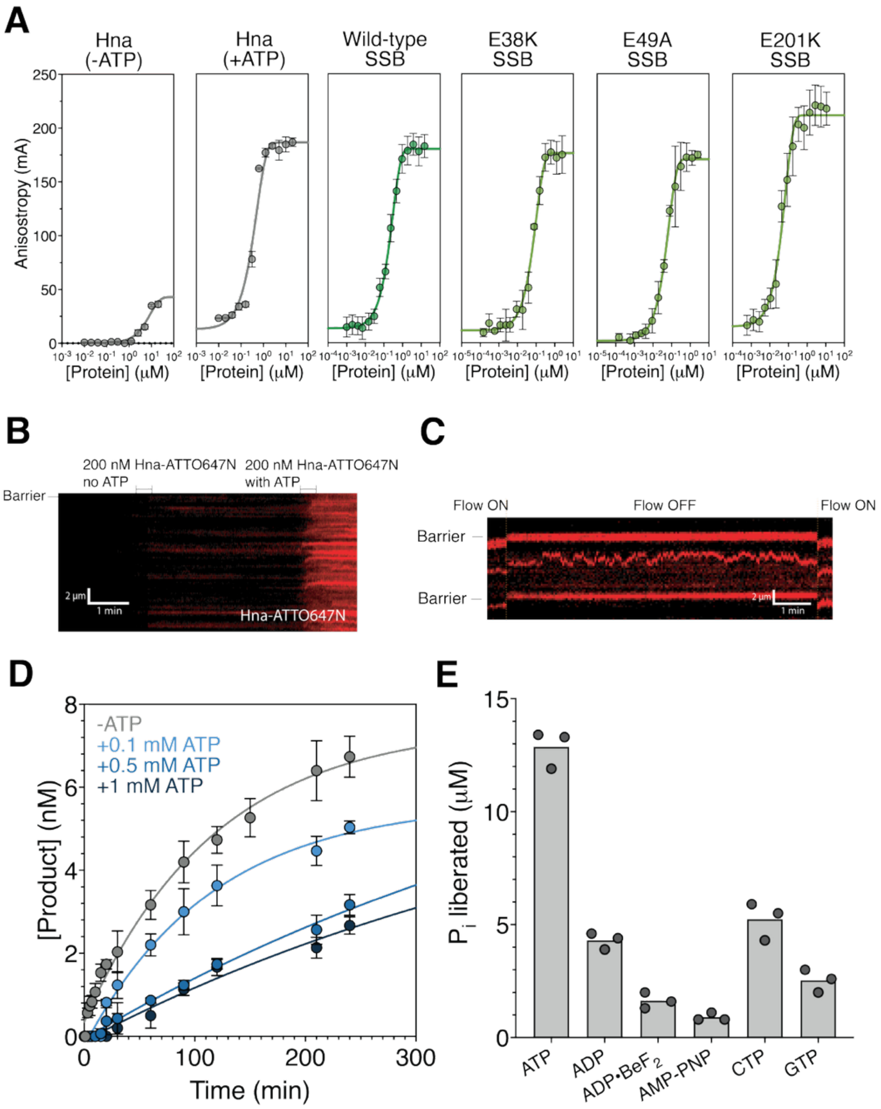


**Extended Fig. 3 | Hna is an ATP-dependent DNA binding protein. A,** Hna and 5A SSB binding curves after incubation with fluorescently labeled single-stranded DNA. Each data point is the average of three independent experiments. **B,** Representative kymograph showing ATP-dependent binding of ssDNA by Hna (red). ATP was injected at the time indicated on the kymograph promoting increased binding events. **C,** Representative kymograph showing passive diffusion of Hna (red) along ssDNA in the absence of buffer flow. **D,** Quantification of DNA cleavage by Hna in the presence of ssDNA with the addition of varying concentrations of ATP using capillary electrophoresis. Data shown are the mean ± standard deviation of three independent experiments for each condition. **E,** Malachite green phosphate detection assay using wild-type Hna with single-stranded DNA present. Phosphate concentration was determined after 30 minutes incubation

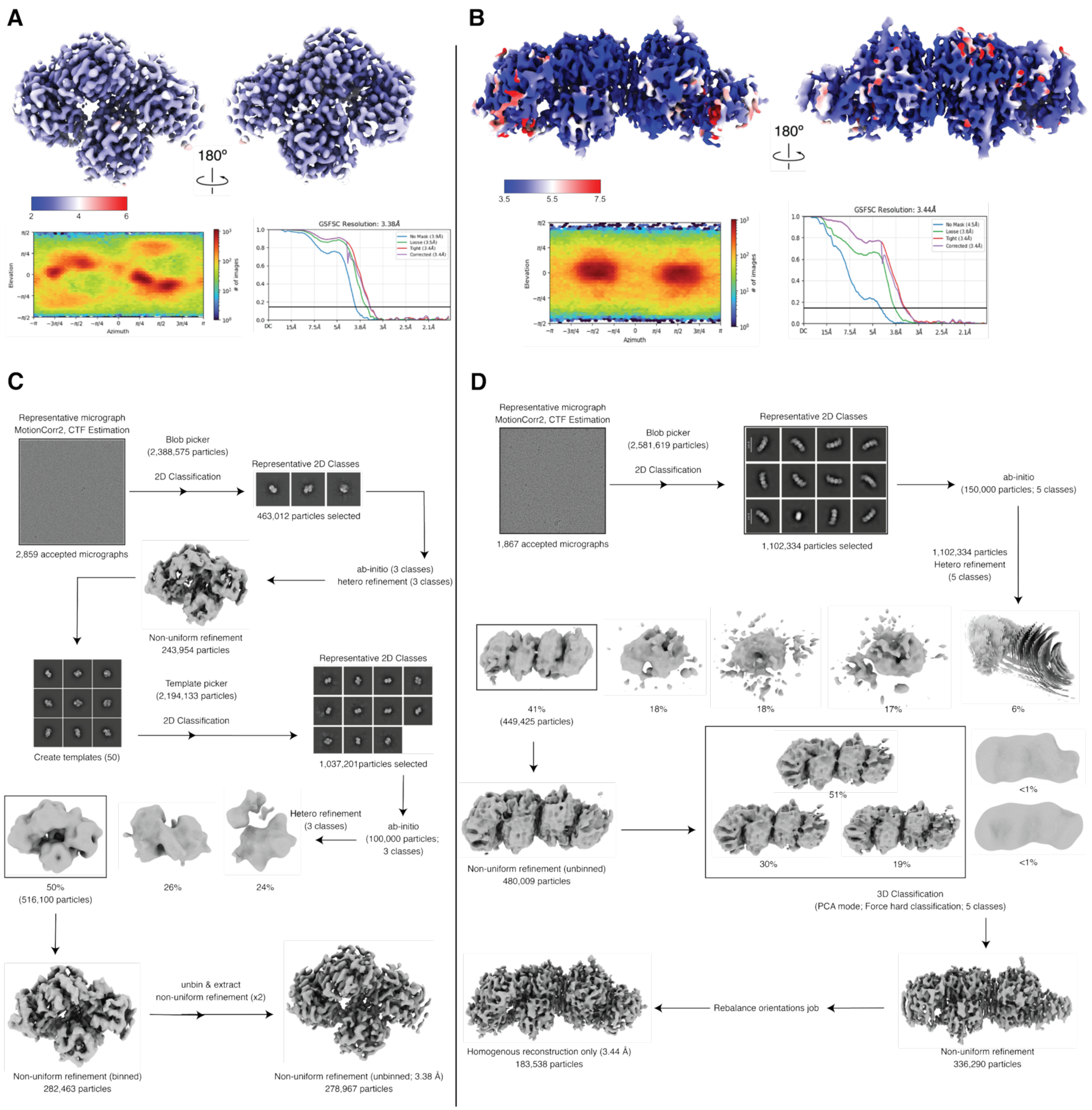


**Extended Data Fig. 4 | Structural analysis of Hna. A,** Unsharpened maps colored by local resolution (top) for Hna monomer and **B,** Hna dimer. Accompanying gold-standard FSC curves for cryo-EM reconstructions (left). with Euler diagrams showing orientation distributions of cryo-EM reconstructions (right). Resolutions were estimated at FSC=0.143. **C,** Data processing pipelines for Hna monomer and **D,** Hna dimer

**Extended Data Table 1 | List of DNA sequences used in the study.**

| Oligo Name | Sequence (5' to 3') |
|---|---|
| ssDNA (55-nt) | agctgacgtttgtatgtctgcgtctcatctttatgcgtcagcagagatttctgct |
| dsDNA complement | agcagaaatctctgctgacgcataaagatgagacgcagacatacaaacgtcagct |
| 3' 1-base overhang complement | gcagaaatctctgctgacgcataaagatgagacgcagacatacaaacgtcagct |
| 3' 3-base overhang complement | agaaatctctgctgacgcataaagatgagacgcagacatacaaacgtcagct |
| 3' 9-base overhang complement | ctctgctgacgcataaagatgagacgcagacatacaaacgtcagct |
| 3' 10-base overhang complement | tctgctgacgcataaagatgagacgcagacatacaaacgtcagct |
| 3' 25-base overhang complement | agatgagacgcagacatacaaacgtcagct |
| 5' 9-base overhang complement | agcagaaatctctgctgacgcataaagatgagacgcagacatacaa |
| Forked complement | tgagacgcagacatacaaacgtcagcttgagacgcagacatacaaacgtcagct |

**Extended Data Table 2 | Cryo-EM data collection, refinement, and validation statistics.**

| | Hna Monomer (EMD-72967; PDB 9YHN) | Hna Dimer (EMD-73047; PDB 9YKJ) |
|---|---|---|
| **Data Collection and Processing** | | |
| Voltage (kV) | 200 | 200 |
| Electron exposure (e-/Å$^2$) | 49 | 49 |
| Defocus range (μm) | -1.5 to -2.5 | -1.5 to -2.5 |
| Pixel size (Å) | 0.94 | 0.94 |
| Symmetry imposed | C1 | C1 |
| Initial particle images (no.) | 2,194,133 | 2,581,619 |
| Final particle images (no.) | 278,967 | 185,538 |
| Map resolution (Å) | 3.3 | 3.4 |
| FSC threshold | 0.143 | 0.143 |
| **Refinement** | | |
| Initial model used (PDB code) | AlphaFold3 | AlphaFold3 |
| Model resolution (Å) | 3.8 | 3.5 |
| FSC threshold | 0.5 | 0.5 |
| Map sharpening B factor (Å$^2$) | 167.3 | 121.9 |
| Model composition | | |
| Non-hydrogen atoms | 6493 | 9547 |
| Protein residues | 821 | 1205 |
| Nucleotides | 0 | 0 |
| Ligands | 3 $Ca^{2+}$ | 2 ADP |
| R.m.s deviations | | |
| Bond lengths (Å) | 0.005 | 0.006 |
| Bond angles (°) | 0.698 | 1.378 |
| Validation | | |
| MolProbity score | 1.12 | 1.99 |
| Clashscore | 2.62 | 6.71 |
| Poor rotamers (%) | 0 | 1.49 |
| Ramachandran plot | | |
| Favored (%) | 97.67 | 91.86 |
| Allowed (%) | 2.33 | 7.46 |
| Disallowed (%) | 0 | 0.68 |
